# Killer Mice: First Documentation of Lethal and Near-Lethal Attacks on Bank Voles by Free-Living Yellow-Necked Mice

**DOI:** 10.64898/2026.04.21.719871

**Authors:** Korneliusz Kurek, Raffaele d’Isa, Michael H. Parsons, Piotr Bebas, Rafal Stryjek

## Abstract

In nature, the most common drivers of lethal aggression are predation and territorial defense. In northeastern Poland, the yellow-necked mouse (*Apodemus flavicollis*) coexists with several rodent species, including the bank vole (*Clethrionomys glareolus*). Compared to voles, *A. flavicollis* is larger, physically stronger, more aggressive, and dominant in the social ecosystem. However, no visually documented instance of a lethal attack by this species has been reported up to date. Here, we present the first recorded case of a fatal attack by a yellow-necked mouse following an encounter with a bank vole. A near-lethal attack is also reported. Importantly, these attacks were not predatory, as no consumption occurred. The attacks appeared instead to be related to interspecies competition, i.e., to competitive interactions between two species that live in the same habitat and use the same type of resources. Notably, while the aggressiveness of yellow-necked mice towards bank voles was known, it was unknown that it could take such extreme forms. Since, in rodents, most competition-related agonistic interactions are aimed at distancing the competitor, the physical destruction of the competitor appears as a surprisingly extreme way of addressing the game of interspecies competition through definitive removal of the opponent. Our observations highlight the need for further research on interspecific aggression among small mammals. They also emphasize the importance of field-based methods, such as camera trapping and continuous video monitoring, which allow for direct observation of animal behavior in natural settings and can reveal rare or previously overlooked interactions.

## 1. Introduction

When different species of rodents inhabit the same environment, and especially when resources are limited, interspecific competition may arise (Grant, 1972; Eccard and Ylönen, 2003). Such a competition may commonly lead to aggressive interactions when interspecies encounters occur (Eccard and Ylönen, 2003). The defended resources may be, for instance, territory, a safe burrow, a food item, or a food cache. Nevertheless, these aggressive interactions among rodents are generally non-lethal, most commonly involving transient confrontations simply aimed at distancing the rival from the defended resource. Although injuries can occasionally occur, reports of fatal outcomes in such encounters are extremely rare.

The yellow-necked mouse (*Apodemus flavicollis*) is well-known for being highly aggressive towards sympatric rodent species (Hoffmeyer, 1973; Montgomery, 1978; Gliwicz, 1981; Grüm and Bujalska, 2000; Mažeikytė 2002; d’Isa et al., 2024a). Yellow-necked mice display this aggressiveness both towards congeners, such as the wood mouse (*Apodemus sylvaticus*) and the black-striped mouse (*Apodemus agrarius*), and towards non-congeners, such as the bank vole (*Clethrionomys glareolus*). Notably, yellow-necked mice are endowed with a physical advantage over all three these sympatric species, being bigger (Górecki, 1965; Barčiová and Macholán, 2006; Bartolommei et al., 2016; Probst and Probst, 2023; Stryjek et al., 2024a) and consequently stronger than all of them, which facilitates their victory in direct fights. In short, yellow-necked mice are characterized by both aggressiveness (tendency to attack) and dominance (ability to subdue) in relation to sympatric rodent species.

For example, laboratory investigations staging encounters in a large indoor pen found that yellow-necked mice attacked and were dominant over wood mice in 79.2% of the cases, whereas wood mice manage to win the fight in just 8.5% cases (Hoffmeyer, 1973). Analogously, when encounters were staged in a testing arena, yellow-necked mice defeated wood mice in 90% of the cases, while wood mice happened to win the fight only in 1.4% of the cases (Montgomery, 1978). Our team, while monitoring free-living yellow-necked mice and black-striped mice in central Poland, observed that, when an interspecific encounter occurred, in 86.7% of the cases it led to agonism (Stryjek et al., 2026a). In 100% of these agonistic interspecific encounters, the yellow-necked mice were the initiator of the agonism. Furthermore, in the vast majority of the cases (84.6%), the yellow-necked mice were dominant^1^ over the black-striped mice.

Bank voles do not only occupy an environment in spatial overlap with yellow-necked mice, but they also share the same microhabitat, as both species are forest-dwellers (Viviano et al., 2022). Bank voles are strongly targeted by yellow-necked mice (Grüm and Bujalska, 2000). Andrzejewski and Olszewski carried out direct observations of the interspecific interactions between wild yellow-necked mice and bank voles, by employing a special observation cabin (elevated 2 m above the ground) located in Poland’s Białowieża National Park (Andrzejewski and Olszewski, 1963). The researchers found that interspecies encounters between yellow-necked mice and bank voles systematically ended with a violent attack by the first and the consequent flight of the second, indicating strong aggressiveness and dominance of the yellow-necked mouse towards the bank vole.

Yellow-necked mice may get to attack voles even if these are already dead at the moment of the encounter, as has been observed in nature, during a field trapping expedition in a lowland forest in southern Moravia (Czechia), when the corpses of the captured animals had been laid out in a row (Zejda, 2002). Notably, in this reported case, the attacking yellow-necked mouse had to pass by 9 dead conspecifics to reach the vole, which was the 10th animal of the row, and it completely ignored all of the conspecifics. Chased away by the human observer, the yellow-necked mouse returned some minutes later, and again it selectively attacked the vole.

Yet, in spite of this well-known aggressiveness, up to now there is actually no report of a yellow-necked mouse in nature having performed a fatal attack against any sympatric rodent species. Remarkably, during our long-term field project studying rodent behavior in central and northeastern Poland (Stryjek et al., 2021a, 2024a; Parsons et al., 2023a; d’Isa et al., 2024a), we observed, in a forest area in northeastern Poland, a unique case in which a free-living *Apodemus flavicollis* individual fatally attacked a free-living *Clethrionomys glareolus*. To our knowledge, this is the first documented instance of such behavior in this species. Additionally, we report a near-lethal attack occurred at a nearby location. This article presents a detailed description of the observed events, along with the ecological and behavioral contexts in which they occurred.

## 2. Materials and Methods

### 2.1 Study site

The recordings were taken in a forest area of Urwitałt (Warmian-Masurian Voivodeship, northeastern Poland), more specifically in a forest-enclosed meadow (53°48′17.00″N; 21°38′28.00″E; altitude: 120 m) near the Lake Łuknajno and belonging to the Masurian Centre for Biodiversity and Education KUMAK, a biological research station of the University of Warsaw.

### 2.2 Animals

The observations were conducted as part of our research project monitoring the behavior of free-ranging rodents (Stryjek and Modlinska, 2016; Modlinska and Stryjek, 2016; Stryjek et al., 2018, 2021a, 2021b, 2024a, 2026a, 2026b; Parsons et al., 2023a; d’Isa et al., 2024a) using the free exploratory paradigm (Parsons et al., 2023b). The subject animals of the present study were free-living individuals of two species, the bank vole (*Clethrionomys glareolus*) and the yellow-necked mouse (*Apodemus flavicollis*), which were observed entering and interacting within test chambers placed in a natural environment. The video recordings, captured from a top-down perspective, did not allow for the determination of the sex of the observed individuals.

### 2.3 Equipment

Two experimental chambers were deployed at the study site. In Figure 1A, the red spot and the blue spot show the locations of chamber 1 and chamber 2, respectively. Both chambers were placed close to the forest’s boundary: chamber 1 was at about 20 m from the treeline and chamber 2 at about 30 m from the treeline. Chamber 1 (external view: Figure 1C; internal view: Figure 1E) had internal dimensions of 37 cm × 27 cm, with a designated observation area measuring 14 cm × 25 cm, which was separated from the rest of the interior by a vertical partition wall. Animals could enter the chamber through a 45 cm long plastic tube with an internal diameter of 4.5 cm, positioned at a 45-degree angle relative to the entrance. Chamber 2 (external view: Figure 1C; internal view: Figure 1E) was 29 cm (length) x 18 cm (width) x 45 cm (height), and contained a transverse partition which divided it into two compartments (northern and southern) connected by a tunnel serving as passage. Both chambers were baited with chocolate-hazelnut cream (Nutella; Ferrero, Alba, Italy) to attract the rodents. The chocolate-hazelnut cream was contained in a circular dish which in chamber 1 was placed in the only accessible compartment, while in chamber 2 was placed in the northern compartment. For both chambers 1 and 2, the entrance/exit of the chamber was always open, so that the animals could access or leave the chambers at any time. In both chambers, video recordings were made employing a Browning Spec Ops Elite HP5 infrared camera, equipped with an additional +4 macro lens to ensure sharp image capture at close range. Recordings were motion-triggered and motion detection was active 24/7.

**Figure 1.**
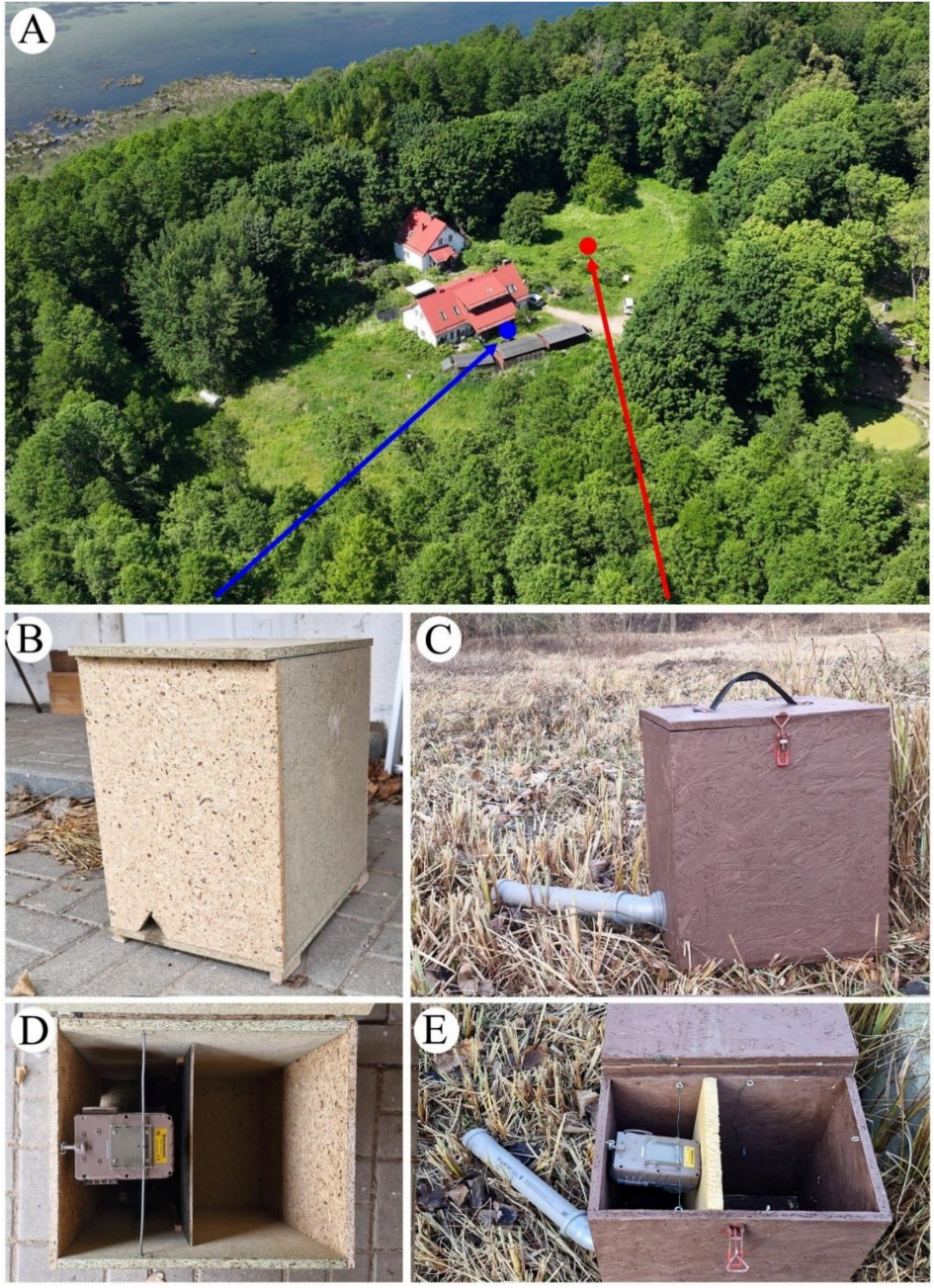
Study site and equipment. A: Observation chambers were deployed in Urwitałt (northeastern Poland), in a forest-enclosed meadow near the Lake Łuknajno, a natural area belonging to the Masurian Centre for Biodiversity and Education KUMAK, a biological research station of the University of Warsaw. The red spot shows the location of chamber 1, while the blue spot shows the location of chamber 2. B and C: external view of chambers 2 and 1, respectively. D and E: internal view of chambers 2 and 1, respectively.

We performed observations from November 2024 to January 2026. More specifically, chamber 1 was deployed on the evening of 18 November 2024 and kept until the evening of 27 December 2024, while chamber 2 was deployed on the afternoon of 19 November 2024 and kept until the afternoon of 7 January 2026. The area where the chambers were deployed features the presence of three major sympatric rodent species: bank voles, yellow-necked mice and black-striped mice^2^ (*Apodemus agrarius*). Within 3 days from deployment of the first chamber, individuals of all three sympatric species began to appear in the chamber. The chamber was deployed at 05:13 PM of 18 November. The first black-striped mouse entered the chamber at 5:40 PM of 18 November, the first yellow-necked mouse at 11:15 PM of 18 November, and the first bank vole at 3:31 PM of 21 November.

### 2.4 Agonism analysis

Encounters between yellow-necked mouse were scored as either agonistic or non-agonistic. An encounter was scored as agonistic either when one of the individuals attacked the other or when one of the individuals fled from the other before being attacked. We then calculated the percentage of agonism as: (number of agonistic encounter/total number of encounters) × 100. Since agonistic and non-agonistic are two mutually exclusive outcomes, the chance level for each of them is 50%. We employed a one-proportion z-test to compare the observed percentage of agonism against the 50% chance level. The software MedCalc v. 23.5.1 (MedCalc Software Ltd, Ostend, Belgium) was employed for statistical analysis.

## 3. Results

### 3.1 Lethal attack by a yellow-necked mouse against a bank vole

The first case we report was observed in chamber 1 on the evening of 15 December 2024, 27 days after the deployment of the chamber. The event occurred at the beginning of the dark phase (encounter: 6:13 PM; vespertine twilight: 3:12 PM; sunset: 3:55 PM), when a bank vole entered the chamber while it was already occupied by a yellow-necked mouse. As this yellow- necked has developed a connection with the chamber when the encounter occurred, before describing the event itself, we report first its background.

On 14 December, at 01:26 PM, a yellow-necked mouse entered the chamber. The mouse paid scarce attention to the food bait (barely eating it), while it appeared more interested in the chamber itself. The mouse was very explorative, moving its head as to sniff/look around, moving about the chamber and rearing to check the camera). Then it got out of the chamber and returned shortly after, resuming exploration, after which it repeated the getting out/getting in/exploring behavioral sequence other 6 times. At 1:55 PM, the mouse started manipulating a few grass blades that were already inside the chamber. In the following hour the mouse was repeatedly getting in and out of the chamber. Around 2:59 PM, the mouse began to bring leaves and grass from outside into the chamber, covering with plant material one of the two dishes containing the cream. It was not possible to say if, in these getting in and out, the mouse was the same, but from the morphological and behavioral consistency it likely was the same.

At 3:19 PM a second yellow-necked mouse appeared in the chamber for the first time, and it was not attacked by the occupant. This second mouse took a few bites of cream and left after about 10 minutes. At 3:49 PM, the first mouse resumed its plant manipulation activity, moving grass and leaves, and making just short breaks for eating. The mouse remained in the chamber alone for the subsequent 5 hours, during which it was either resting or nest-building^3^. At 8:50 PM a second yellow-necked mouse returned, remained inside for about 15 minutes and then left. This second mouse was not attacked by the first yellow-necked mouse. At 9:07 PM, the first mouse started going out to find plant material and carry it inside the chamber. The plant-carrying and nest-forming went over for the following 2 hours. At 11:10 PM the mouse started covering the second dish of cream, which at 11:40 PM was totally covered, by using the plant material in the chamber plus other material carried from outside. Then the mouse began to cut with its teeth some long grass blades and positioning them around the box. At 0:15 AM of 16 December, the mouse ceased its nest-building activity. Between 1:22 AM and 5 AM, the mouse was resting or sleeping. At 5:16 AM it resumed nest-building, until 6:07 AM, and at 6:46 AM it left the chamber.

Between 7:19 AM and 7:31 AM, the chamber was visited by a black-striped mouse, which ate some cream. Shortly after, a bank vole visited the chamber from 7:37 AM to 7:40 AM, eating cream as well. Between 7:45 AM and 3:30 PM, there were several visits of either black-striped mice or bank voles, which showed regular foraging activity. At 3:31 PM, when a black-striped mouse was inside, a yellow-necked mouse returned, likely the same one that had spent the previous afternoon, evening, night and early morning nest-building and resting in the chamber^4^. The yellow-necked mouse immediately attacked the black-striped mouse, which first hid in the grass material and soon after sneaked out of the chamber. At 3:40 PM the yellow-necked mouse left and at 4:05 PM it returned carrying inside more plant material. After that the yellow-necked mouse went on bringing inside plant material, at 4:14 PM a black-striped mouse entered and got immediately attacked by the yellow-necked mouse, in response to which it promptly escaped from the chamber. After this, the yellow-necked mouse continued its nest-building activity. From 5:48 PM to 6:08 PM, the mouse was modelling the nest to give a rounded shape with a central resting space.

Finally, at 6:13 PM the fatal encounter between the bank vole and the yellow-necked mouse occurred. The ensuing attack was very violent and lasted approximately 20 seconds, with only short interruptions, leaving the bank vole agonizing. The detailed dynamics of the attack is shown in Figure 2 and Supplementary Video 1. Basically, the yellow-necked mouse occupying the chamber was engaged in nest-building (Figure 2A). The vole entered the chamber and it immediately got aggressively chased by the mouse (Figure 2B). The vole managed to hide in the south-eastern corner of the chamber and the mouse started looking for the vole inside the tube (Figure 2C), eventually exiting for 4 seconds, during which the vole at first hid on top of the tube (Figure 2D) and then went back to the same corner of the chamber as before. When the mouse returned, it detected the vole in the corner (Figure 2E) and attacked it (Figure 2F), after which a very intense full-contact fight occurred, in which the vole was thrown about the chamber for a couple of seconds, until it was able to escape once more in the south-eastern corner (Figure 2G: the signs of the bites are visible on the vole’s dorsum). Once again, the mouse looked for the vole inside the tube (this time for less than a second) and in the meanwhile the vole hid on the tube. Not finding the vole inside the tube, the mouse resumed looking for it in the chamber. In less than a second, the mouse found the vole and a 3-second chase started, after which the vole was caught and a very convulsive fight of about 8 seconds followed (Figures 2H and 2I). This fight left the vole severely injured, lying prone in the south-eastern corner, apparently incapable of fighting back or escaping.

**Figure 2.**
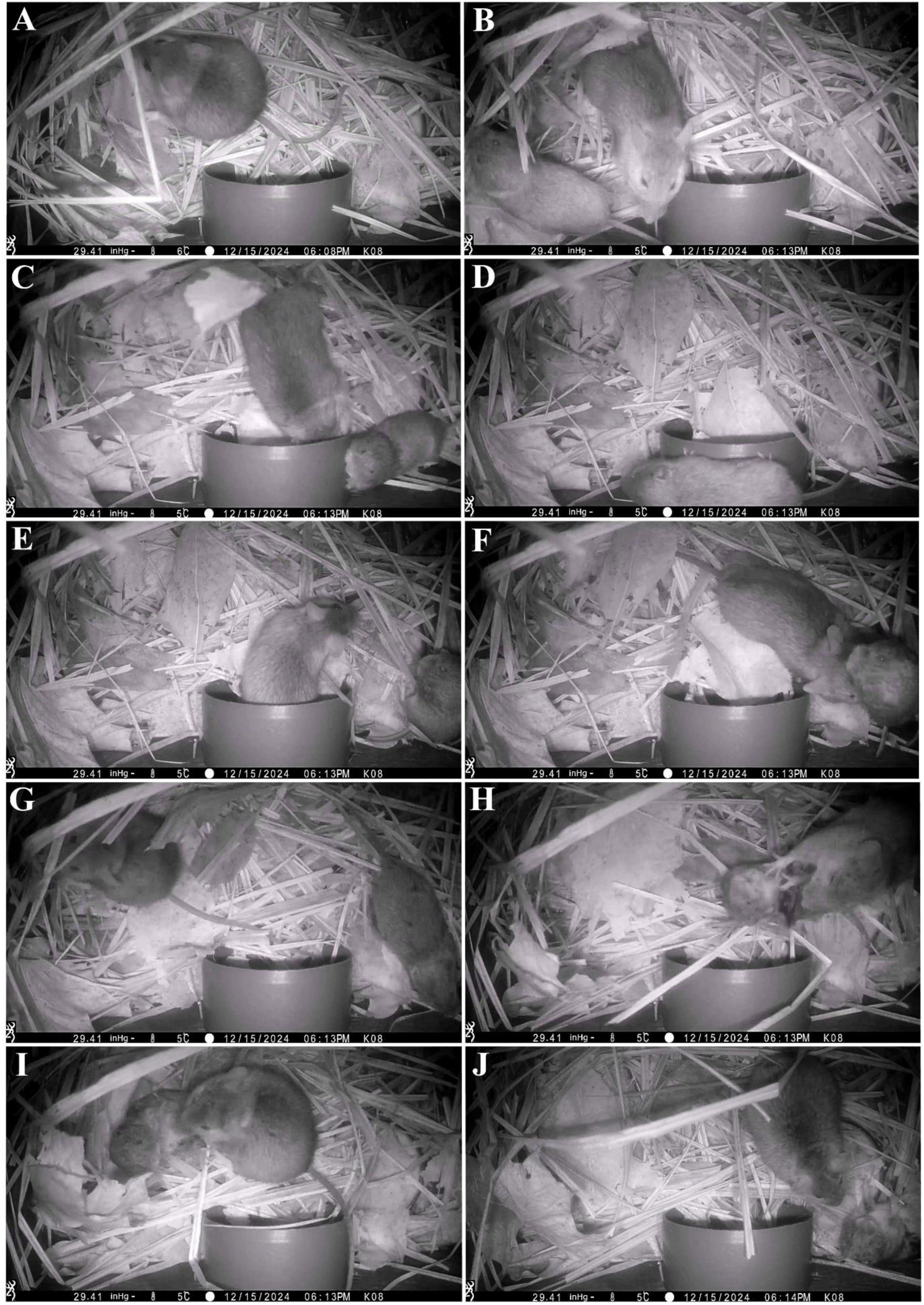
Lethal attack by a yellow-necked mouse towards a bank vole.

Strikingly, although at this stage the vole was clearly neutralized, the mouse struck the moribund vole another three times, demonstrating an extremely high aggressiveness. The vole did not react to the first strike, while after the second strike it just turned, lying with the back on the ground (Figure 2J). At this point, the vole was showing clear signs of agony (i.e., lateral recumbency, convulsions, rhythmic gasping, and absence of defensive fighting against the aggressor). Nevertheless, the mouse attacked again a third time, with no reaction by the vole. Within 4 minutes from the mouse’s last strike, the vole stopped gasping and any visible respiration ceased entirely, indicating that it had died.

Notably, during the confrontation, the vole had taken several times the submissive posture (lying on the back, exposing the ventral vulnerable body parts such as abdomen and throat, and raising the forelimbs). In rodent agonistic^5^ encounters, submissive postures (including the supine on-back posture and the submissive upright posture) generally inhibit further bites from the attacker (Scott and Fredericson, 1951; Grant, 1963; Grant and Mackintosh, 1963; Blanchard et al., 1977; Koolhaas et al., 1980; Miczek et al., 2011; Battivelli et al., 2024; Zhukov et al., 2024). Nevertheless, the yellow-necked mouse was not stopped by this. When, at the end of the confrontation, the vole was left agonizing, it was in a submissive posture, which it maintained for the whole agony period. Notwithstanding this, the yellow-necked mouse newly attacked the vole, which did not change posture in spite of the new aggression and died in the submissive position.

At 7:30 PM a second yellow-necked mouse entered the chamber. The first mouse immediately approached the newcomer, which initially retracted as if scared from the sudden approach in the darkness, but then the first mouse started sniffing the second mouse (on the head, the anogenital area and the tail), without biting the second mouse, and it accepted the newcomer’s presence in the chamber. No fight occurred. On the contrary, they showed high sociability one to the other, both sniffing the other and touching with their forepaws the back of the other (Figures 3A and 3B). Then the two mice huddled together in the nest, mutually seeking physical contact, and alternating rest with sniffing and grooming of the other (Figures 3C-3F; Supplementary Video 2). At 10:30 PM, one of the two yellow-necked mice (likely the second mouse) left the chamber.

**Figure 3.**
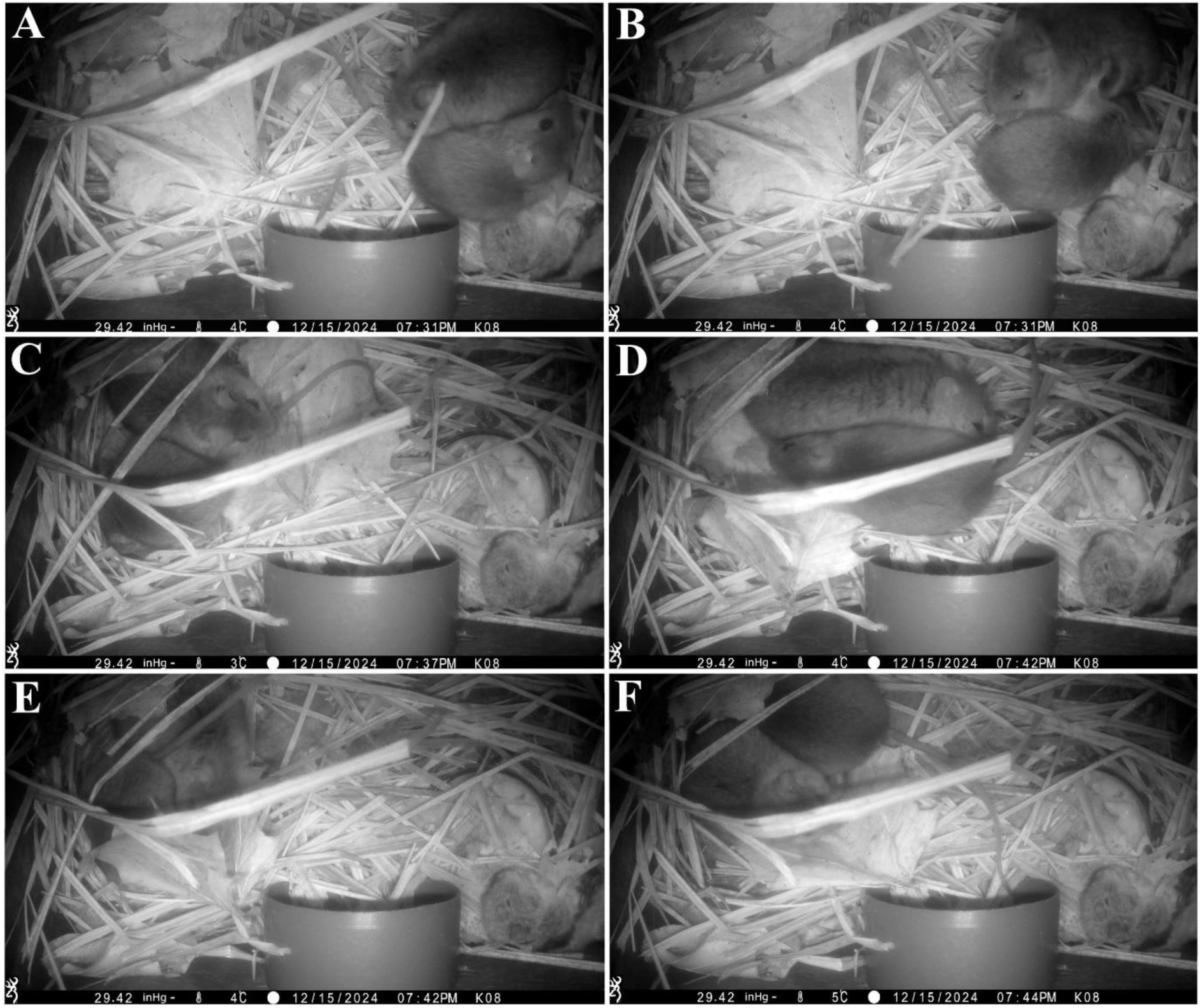
Aftermath of the lethal attack by a yellow-necked mouse towards a bank vole.

After this, at 11:38 PM, the remaining mouse began nest-building. Within this activity, the first operation performed by the mouse, between 11:38 PM and 11:44 PM, was to bring inside some leaves and specifically use them to cover the body of the dead vole (Supplementary Video 3). Then the mouse started covering with leaves the whole floor of the chamber, continuing with great persistence and finishing this nest-building activity only at 5:19 AM. While the mouse was nest-building, at 3:22 AM, a second yellow-necked mouse came again inside the chamber, for about one minute. As previously, the incomer was not attacked. The two conspecific mice showed a high sociability one to the other, with abundant mutual social investigation featuring sniffing and touching each other with the forepaws. About one hour after having ceased nest-building, the mouse left the chamber (at 6:31 AM). Remarkably, none of the two yellow-necked mice showed interest for consumption of the dead vole, which the following afternoon was found by the experimenters unconsumed. Over the course of the entire period of observation, no other encounter occurred in chamber 1 between yellow-necked mice and bank voles, neither before nor after the lethal encounter.

### 3.2 Near-lethal attack by a yellow-necked mouse against a bank vole

The second case occurred on the morning of 12 October 2025, few minutes after dawn (encounter: 7:04 AM; sunrise: 6:55 AM), when a bank vole already in the chamber underwent a near-lethal attack by an intruding yellow-necked mouse. In this case, none of the involved individuals had been previously nesting in the chamber. This case can be subdivided into two contiguous episodes, featuring escalating levels of aggressiveness. The first episode is depicted in Figure 4 (the full scene can be seen in Supplementary Video 4).

**Figure 4.**
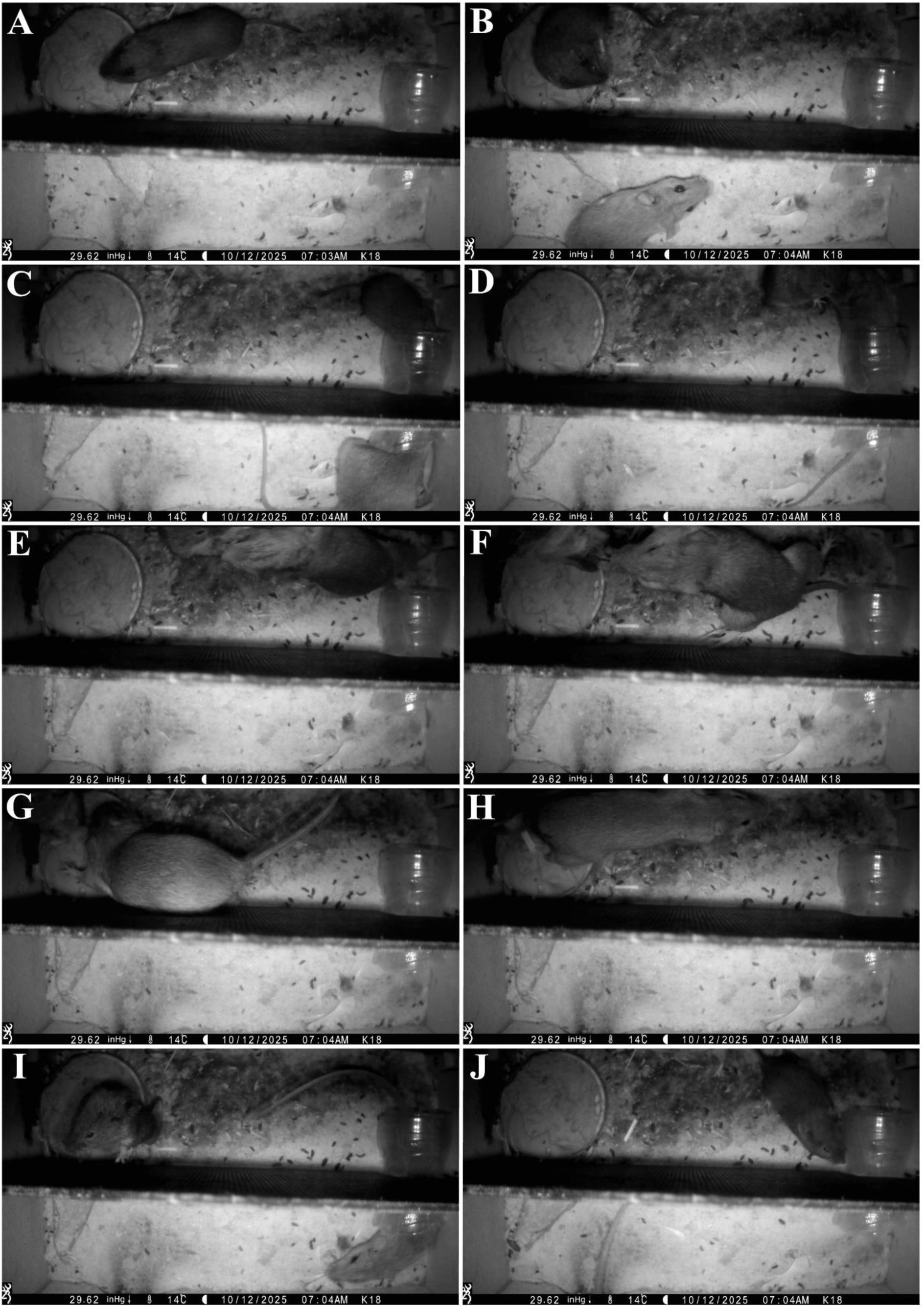
Near-lethal attack by a yellow-necked mouse towards a bank vole (episode 1)

At 7:03 AM, a bank vole entered the chamber, reached the plate containing the food and started eating (Figure 4A). At 7:04 AM, a yellow-necked mouse entered the chamber still occupied by the vole (Figure 4B). Detecting the arrival of another animal, the vole turned towards the entrance tube (Figure 4B) and moved to the tube, putting the head inside it to gain cues about the newcomer (Figure 4C). At the same time, the mouse reached the tube as well (Figure 4C) and, upon making contact with the vole, it immediately attacked it (Figure 4D). The mouse attacked so vigorously that it threw the vole into the north-western corner (Figures 4E-4G). Then the mouse turned (4H), returned to the tube and crossed it leaving the vole in the north-western corner (Figure 4I). Once in the southern section of the chamber, the mouse exited the chamber and the vole, now alone, moved next to the tube remaining in vigilant monitoring (Figure 4J). The fight lasted about a second and the mouse’s permanence time in the chamber was about 2 seconds. After the attack, the mouse showed no interest for the food, the vole or the chamber itself.

The second episode (Figure 5; Supplementary Video 4) occurred immediately after the first. Four seconds after leaving the chamber, the mouse came back (Figure 5A). Again, the mouse quickly crossed tube, reached the vole in the northern section and attacked it (Figure 5B). A very violent fight followed, during which the vole was repeatedly bitten (Figure 5C). After about 8 seconds of fight, the vole took the on-back submissive posture next the tube (Figure 5D) and the mouse paused the attack. Nevertheless, after 3 seconds, when the mouse tried to move away, the mouse resumed the attack and repeatedly bit the vole with even greater violence (Figure 5E and 5F). The new fight lasted about 4 seconds. This time the mouse left the vole very severely wounded and disabled (Figure 5G). After having neutralized the vole, the mouse crossed the tube to reach the southern section (Figure 5H) and left the chamber (Figure 5I). Like in the first episode, after attacking the vole the mouse exhibited no interest for the food, the vole or the chamber. While the mouse was leaving, the vole started crawling and crossed the tube as well (Figure 5H). After the mouse had left, the vole followed it (Figure 5I) and exited the chamber too (Figure 5J). The vole was severely impaired in its movement and could not lift its right hindlimb, which was visibly dragged (Figure 5I and Supplementary Video 4). Out of the chamber, it is unknown how much the vole could have survived after these wounds. However, even if it did not die in the immediate aftermath, in this state of disability its chances of survival were significantly reduced due to reduced capacity to forage and escape predators. It is therefore almost certain that this attack caused a premature death of the vole.

**Figure 5.**
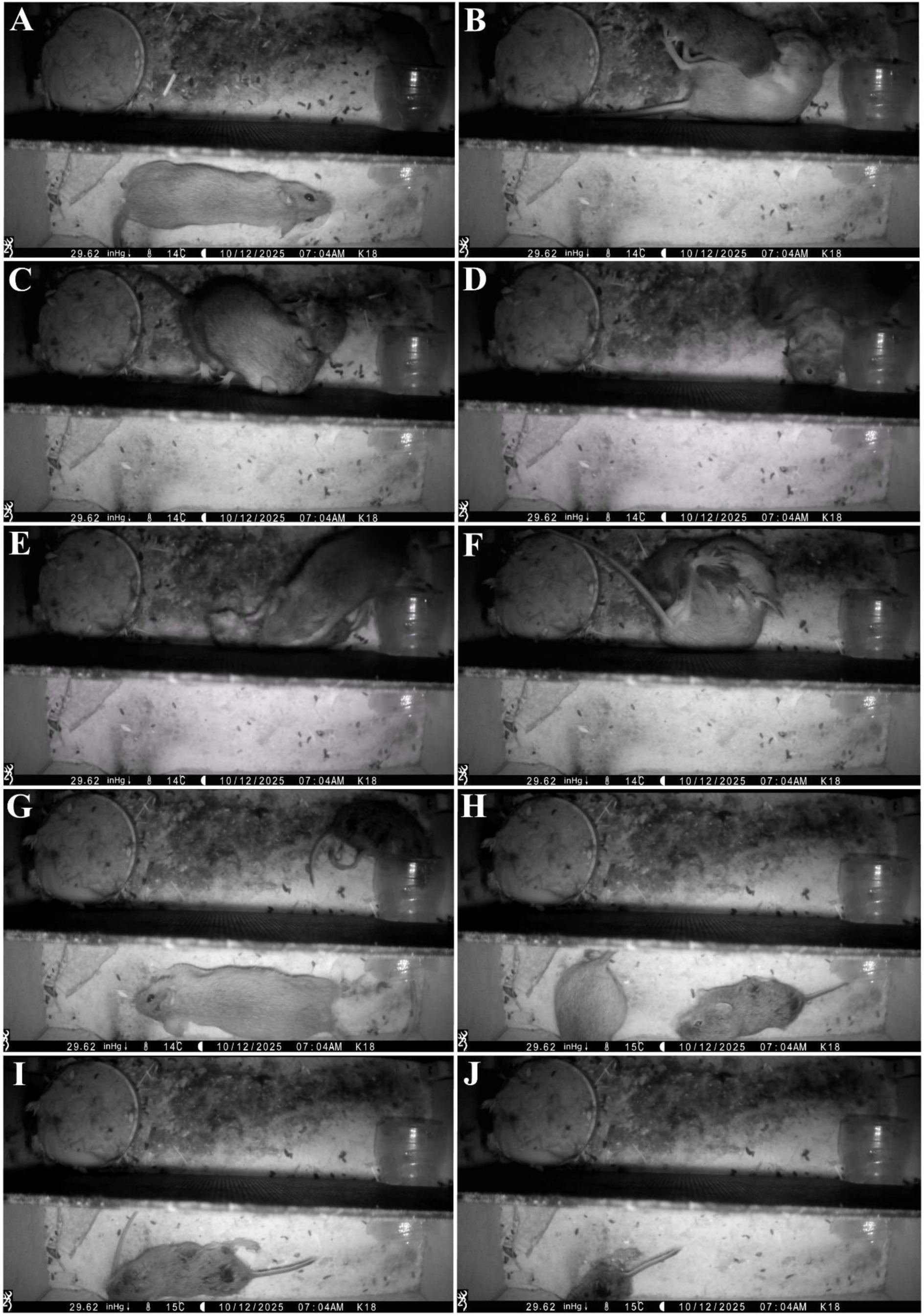
Near-lethal attack by a yellow-necked mouse towards a bank vole (episode 2)

In chamber 2, over the course of the entire period of observation, we observed in total 15 encounters between yellow-necked mice and bank voles (the near-lethal case plus another 14). Notably, in these encounters, agonism was observed in 100% of the cases. Considering agonistic and non-agonistic as two mutually exclusive outcomes, the percentage of agonism in the encounters between yellow necked mice and bank voles was significantly over chance level (One-proportion z-test: z = 3.873; p = 0.0001). Among the agonistic encounters between yellow-necked mice and bank voles, in 66.7% of the case the vole escaped before being attacked, while in the remaining 33.3% of the cases the vole left the chamber after having been attacked.

## 4. Discussion

The most striking aspect of the two described cases is the level of aggressiveness that the yellow-necked mice displayed against the bank voles. In both cases, when the encounter occurred, the yellow-necked mouse attacked the bank immediately, reacting with extreme swiftness to the presence of the vole. Such promptness raises the question of which sensorial cue could have triggered the attack. In case 1, the killer mouse immediately attacked an incoming vole, but it never attacked when a conspecific entered. How could the yellow-necked mouse discriminate between a conspecific and a vole? Interestingly, in the chamber there was complete darkness, so it is possible to rule out that the identification of the target could have been based on visual features. An alternative possibility is that the yellow-necked mouse could have distinguished an incoming conspecific mouse from a vole just by the sound of their walking. However, in the lethal case, the floor of the chamber, as well as the floor of the exit of the tube, were covered with soft nesting material, so this possibility seems unlikely.

The most plausible hypothesis is that the yellow-necked mouse was reacting to olfactory cues, and that the odor of a vole triggers a prompt and strong attacking behavior in yellow-necked mice. This hypothesis is consistent with the behavior of the attacker in the case described by Jan Zejda, where a yellow-necked-mouse repeatedly attacked a dead vole (Zejda, 2002). In this case, 16 dead rodents had been laid on the ground in an oval meadow which was completely surrounded by forest. Only 2 of the dead rodents were bank voles (the 10th and the 12th), while the other 14 bodies belonged to yellow-necked mice. At a certain point, a large yellow-necked mouse emerged from the forest underground from the side at the left of the row of corpses. It moved towards the dead bodies, but it passed by 9 conspecifics, without even sniffing any of them, and attacked the 10th animal, which was the vole closest to it. Chased away, the mouse ran into the forest at the north of the row of bodies. Later on, when the yellow-necked mouse returned, it came from the northern side and this time it moved perpendicularly towards the corpses, choosing a trajectory that reached directly the 10th animal (the vole) without passing by any other animal, and very quickly attacking and dragging the vole. Zeyda hypothesized that the persistently selective attack of the yellow-necked mouse could have been triggered either by the different coat color of the vole or by its odor.

Moreover, in our observed cases, in addition to the immediacy, another notable characteristic of the attacking behavior of the yellow-necked mice is its exceptional relentlessness. Indeed, in both cases, the voles repeatedly took the supine submissive posture. Figure 6 shows close-ups from the lethal case (Figure 6A) and the near-lethal case (Figure 6B). Lying on the back (the supine posture) is a typical sign of submission present in many rodent species including voles, such as the field vole *Microtus agrestis* (Clarke, 1956) and the bank vole (Kruczek and Styrna, 2009). In rodents, generally, when an individual takes the submissive posture after having being attacked, this reduces the likelihood of further attacks, often leading to cessation of aggression by the dominant individual (Scott and Fredericson, 1951; Grant, 1963; Grant and Mackintosh, 1963; Blanchard et al., 1977; Miczek et al., 2011; Battivelli et al., 2024; Zhukov et al., 2024). Nevertheless, we observed that, in both the lethal and the near-lethal cases, the yellow-necked mice continued to attack in spite of the target’s submission. Moreover, in the lethal case, even if the major fight had left the vole severely injured, neutralized (incapable of fighting) and immobile in a corner, this did not stop the yellow-necked mouse from launching other 3 attacks against the vole, which did not fight back any more. In the second attack, the vole just took the submissive supine posture. After that, the target was agonizing and essentially unresponsive. Nonetheless, the yellow-necked mouse resumed attacking. Overall, the behavior of the yellow-necked mouse appears to be an over-overkilling. Remarkably, in rodents, attacking notwithstanding the opponent’s submissive posture and regardless of its unconsciousness have been identified as two main criteria to distinguish ordinary aggression from aberrant aggression, which is technically defined as violence (Natarajan et al., 2009, 2010). In our observed cases, the behavior of the yellow-necked mouse against the vole displays the features of violence rather than those of basic aggression. The attacking behavior of the yellow-necked mice, with its incapacity of being modulated by either submission or physical unresponsiveness (agony) of the opponent, denotes inflexibility, and consequently violence.

**Figure 6.**
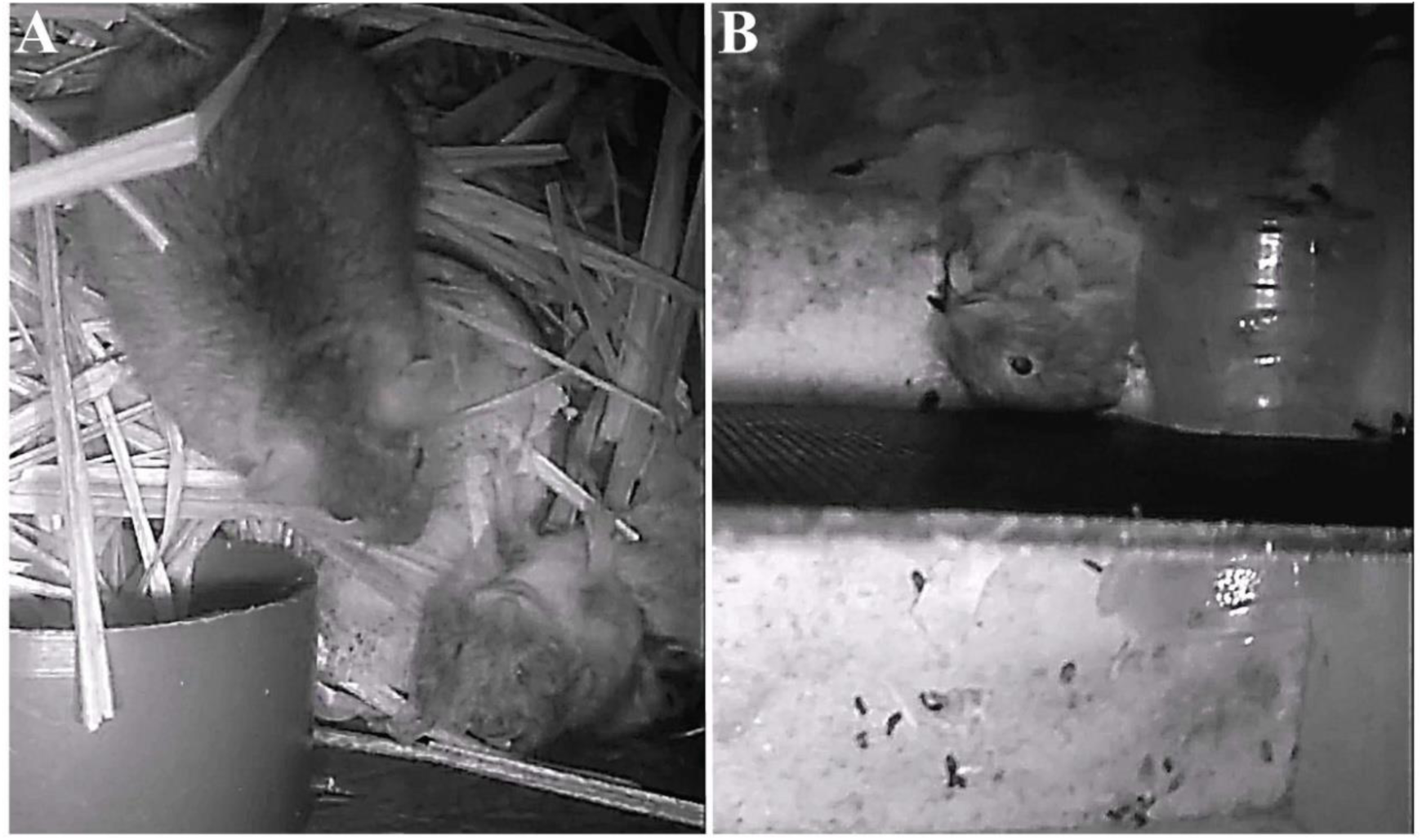
Close-ups of vole supine submissive postures in the lethal (A) and near-lethal (B) encounter.

Finally, another prominent aspect of the yellow-necked mouse’s aggressiveness is the intensity of the attacks (i.e., the amount of damage inflicted). In one case the attack was lethal and in the other it was near-lethal (but most likely it had lethal consequences). In the lethal case, the persistent attacking of an agonizing vole was not just qualitatively aberrant, but also showed a quantitively exaggerated amount of damage inflicted.

In sum, the immediacy, relentlessness and intensity of the yellow-necked mice’s attacking behavior were all extreme. Why were the yellow-necked mice so aggressive against the voles? Remarkably, we can be sure that the yellow-necked mice’s attacks did not have predatory scopes, as in both cases the attackers completely ignored the vole as food source. In the lethal case, the killer remained for hours in the chamber and did not even attempt to consume the vole’s body. In the near-lethal case, the attacker left the chamber immediately after having neutralized the vole. We can therefore be reasonably sure that the yellow-necked mice were not predating (i.e., trying to obtain meat to eat).

It appears instead that the vole-attacking behavior of the yellow-necked mice was related to interspecies competition rather than to predation. Remarkably, competition-related aggressiveness can be of two types: contingent (over specific resources that are present at that moment, such a food item) or unconditional. While the first type is driven by the individual goals of the attacking animal (obtaining a specific and contingent resource), the second type is innate and is provided by natural selection to an entire species to oppose another species as a whole. Regarding our observed cases, were the attacking yellow-necked mice driven by contingent competition-related aggressiveness or by unconditional competition-related aggressiveness? In order to reply to this question, we should consider over which contingent resources the yellow-necked mice could possibly have been fighting (i.e., what specific resource could they have obtained by eliminating the vole). In the observed situations, there seem to be two possible resources: the food (the chocolate cream) and the chamber itself (which would represent a shelter from natural elements and predators). Were the yellow-necked mice fighting for these resources?

In the lethal case, it is possible that the yellow-necked mouse’s aggressiveness towards the vole was further exacerbated by the fact that the mouse perceived the vole’s entry as a nest-invasion. Indeed, just before the fatal event, on the day of the killing and on the previous day, the killer mouse had been persistently nest-building and it had occupied the chamber for a very long period of time. On the day of the killing, after having spent the whole night inside the chamber, the yellow-necked mouse left it around 7:00 AM and returned at about 15:30 PM. Yellow-necked mice are highly nocturnal, with a percentage of nocturnality (i.e., percentage of activity during the dark phase) up to 95% (Stryjek et al., 2026a). In a previous study of ours, where we analyzed the temporal preferences of yellow-necked mice for foraging, we saw that they were active mainly between 18 PM and 7 AM, while between 7 AM and 17 PM their activity was almost null (Stryjek et al., 2026a). On the day of the killing, the time window during which the yellow-necked mouse was away from the chamber (7 AM to 15.30 PM) corresponds to the time window when yellow-necked generally cease to show extra-burrow activity as they have retreated to their primary nest to sleep. The fact that the yellow-necked mouse left the chamber exactly for that time window suggests that it was not considering the chamber as its primary nest. Nevertheless, its prolonged nest-building activity and occupation of chamber suggest that the chamber could have been for the yellow-necked mouse either a secondary nest or a potentially new primary nest in the future. In both cases, for the killer mouse the chamber would have been a valuable resource to defend. Therefore, the motivation to defend a contingent resource (the chamber) could have intensified the mouse’s aggressiveness. Another factor possibly increasing the perceived value of the chamber as a resource to defend, and consequently the aggressiveness of the yellow-necked mouse, could be the low temperature. In the night preceding the lethal attack, the temperature has dropped subzero (to -1 °C). On the other hand, in the night preceding the near-lethal case, the minimal temperature had been 10 °C.

However, even in the other observed vole-attack (the near-lethal case), the aggressiveness of the yellow-necked mouse was still very high, and in that case the mouse was clearly not defending any contingent resource, since it entered, neutralized the vole and came out, all within just a few tens of seconds, ignoring both the food (the chocolate cream) and the chamber itself. The attacker did not appear to have any interest for either the cream or the chamber. On the other hand, the context of the yellow-necked mouse’s entry suggests that the attacker was interested in the vole itself rather than in any contingent resource. Indeed, the yellow-necked mouse entered the chamber almost together with the vole, just 7 seconds after it, a time so short that it is unlikely to be a coincidence and which suggests that the yellow-necked mouse was actually following the vole from the start. After having neutralized the vole, the mouse immediately left the chamber, without even approaching the dish with the cream.

Altogether, these facts suggest that yellow-necked mice may be endowed with an instinctual attraction for the odor of weaker sympatric rodents, along with an instinct to violently attack them. A study explored the reactions of yellow-necked mice and common dormice (*Muscardinus avellanarius*) to the odors of the nests of sympatric rodents by employing a two-choice test (Zaytseva-Anciferova and Nowakowski, 2012). Basically, wild-caught individuals were released in a T-shaped apparatus where at the end of each arm there was a chamber. One chamber contained an odor (nest material of a given species), while the other was empty. The animals could freely choose where to enter. If the animals chose to enter the chamber with the nest odor, the test was scored as positive. Common dormice showed a strong avoidance of the odor of yellow-necked mice (just 35% of positive tests). In contrast, yellow- necked mice displayed a much higher percentage of positives for the odor of dormice, both for that of the sympatric common dormouse (59%) and for that of the allopatric woodland dormouse *Graphiurus murinus* (67%), which belongs to the family of Gliridae like the common dormouse but could not have been encountered previously by the yellow-necked mice. Towards the odor of conspecifics, yellow-necked mice showed only 38% of positives.

Notably, there is evidence suggesting that yellow-necked mice enter the nests and kill the young of common dormice (Juškaitis, 2008, 2014; Zaytseva-Anciferova and Nowakowski, 2012). Common dormice are arboreal rodents that move, forage and sleep on trees (Juškaitis and Büchner, 2013; Gubert et al., 2023). Yellow-necked mice are proficient tree climbers (Borowski, 1962; Montgomery, 1980; Marsh, 1999) and, in addition to underground burrows, they employ tree nests as well (Truszkowski, 1974; Juskaitis, 1997; Marsh, 1999). In Lithuania, if in autumn arboreal nestboxes are left in the forest, yellow-necked mice generally occupy 10-13% of them (Juškaitis, 2000). In autumn 2007, 6 young common dormice were found killed in the nestboxes, supposedly by yellow-necked mice (Juškaitis, 2014). Several pieces of evidence pointed to this conclusion. First, that autumn yellow-necked mice were particularly abundant and their percentage of nestbox occupation was extraordinarily high, reaching 35%. Second, three dormice were found killed in a nestbox which, after removal of the corpses, was seen being occupied by yellow-necked mice. Third, two of the nestboxes which contained killed dormice had been filled with hazelnuts, a food item which is specifically hoarded by yellow-necked mice. Common dormice do not cache hazelnuts. Rather, they eat hazelnuts in the tree canopy where they find them. Interestingly, the bodies of the dormice had sign of bites, but had not been consumed, indicating that predation was not the prime motivation of their killing. Consequently, it appears that these killings were related to interspecies competition. Indeed, the yellow-necked mouse’s tendency to enter the nest of dormice and kill them would explain both the common dormouse’s avoidance of the yellow-necked mouse’s odor and the yellow-necked mouse’s attraction for the common dormouse’s odor. While more investigations are necessary to understand if yellow-necked mice are actually attracted by the odors of weaker sympatric rodents, this hypothesis should be considered in future research on the interspecies interactions between these species.

A last notable aspect regarding the yellow-necked mouse is the long-term behavior of the killer mouse towards the corpse of the killed vole. As previously highlighted, the killer did not consume the vole after killing it. Most interestingly, at about 5.5 hours from the vole’s death, the mouse started bringing inside the chamber some leaves and specifically used them to cover the corpse of the vole, operation which was completed in 6 minutes. This corpse-burying behavior could be related to the fact that decaying bodies produce decomposition-related organic compounds that contribute to the so-called odor mortis, such as putrescine and cadaverine, two substances characterized by a highly unpleasant odor. Decomposition starts 4 minutes after death and, during this process, the production of decomposition-related odorous compounds causes an increasingly strong odor mortis (Cieśla et al., 2023). Considering the sensitivity of their olfaction, rodents could be easily disturbed by decomposition-related odors. Indeed, laboratory research has shown that rats bury the bodies of dead conspecifics (Pinel et al., 1981; Gonçalves and Biro, 2018). However, they buried the bodies of conspecifics that had been dead for more than 40 hours, but not the bodies of conspecifics that had been dead for less than 5 hours (Pinel et al., 1981). Further experiments showed that this corpse-burial behavior is elicited by putrescine and cadaverine. Indeed, it was seen that if wooden dowels were soaked with either putrescine and cadaverine, in both cases they were buried by the rats (Pinel et al., 1981). Impressively, if live but anesthetized conspecifics were sprinkled with putrescine or cadaverine, they were buried in the same manner (Pinel et al., 1981). In contrast, if rats had been rendered anosmic, then they did not bury aged carcasses nor wooden dowels sprinkled with putrescine or cadaverine (Pinel et al., 1981). In our case, the external temperature at the time of the vole’s death was low (2 °C). However, the presence of punctures in the flesh due to the mouse’s bites could have made some decomposition-related odors detectable by the mouse already at 5.5 hours post-mortem. Additionally, it is possible that during its agony the vole lost control over its bladder and/or anal sphincter, releasing urine and feces that may have contributed to create an odor perceived as disturbing by the yellow-necked mouse.

While, up to here, we have discussed mainly the behavior of the attacking species, it is important to make some considerations about the behavior of the attacked species as well. For what concerns the voles, the most notable aspect of their behavior was their helplessness upon being attacked by a yellow-necked mouse. Within comparative psychology (d’Isa and Abramson, 2023), the comparison of the behaviors of two closely related species in the same context can help to better identify the species-specific behavioral peculiarities of each species. On the same day of the vole-killing, we observed, in the same chamber, two cases of attack by the yellow-necked mouse upon the black-striped mouse. In both cases, the black-striped mouse rapidly evaded the yellow-necked mouse and escaped from the chamber, avoiding fatal consequences or severe injuries. In a previous work of ours, we saw that, while in interspecies encounters the yellow-necked mice behaved very aggressive towards black-striped mice (generally them attacking within 2 seconds from the beginning of the encounter), in the vast majority (84.6%) of these agonistic encounters the black-striped mice promptly left the chamber, and rarely (in 15.4% of the cases) they managed to expel the attacker (Stryjek et al., 2026a).

On the other hand, in our two reported cases, when the bank voles were attacked, they did not take the tube to escape. Both voles surely knew the position and the function of the exit tube, as they had just used it to enter. Nevertheless, it appears that, after being attacked so violently by a yellow-necked mouse, the bank voles likely entered a panic-like state that impeded them to carry out an effective decision making which could have led them to a successful escape. Over the course of the whole period of our observations, the total number of agonistic interactions between bank voles and yellow-necked mice (in either chamber 1 or 2) was 16. Within these agonistic encounters, in most cases (62.5%) the voles fled before any contact with the yellow-necked mouse, and escaped uninjured from the chamber. On the other side, if the vole waited for the yellow-necked mouse’s first strike, the voles were neutralized or killed in 33.3% of the attacks by yellow-necked mice. As we observed in a previous study of ours (Stryjek et al., 2026a), for the black-striped attacked by yellow-necked mice, this percentage was instead zero. It appears that bank voles are particularly disadvantaged in the case of a fight with a yellow-necked mouse and that, in the case of an encounter with this species, their best chance of survival is immediate fleeing.

While yellow-necked have been previously presumed to kill sympatric rodents through indirect evidence, the present article is the first work showing direct evidence of this. For the first time, we here show photographic and videographic evidence of vole-killing by yellow-necked mice. How often could voles be killed by yellow-necked mice in nature? Why have these killing not have been documented before?

We propose that similar incidents may occur in nature more frequently than currently recognized. *Apodemus flavicollis* and *Clethrionomys glareolus* are sympatric species occupying similar ecological niches (forest areas) and often competing for food or spatial resources such as burrows or nesting chambers. Direct, visual observation of encounters between bank voles and yellow-necked mice (Andrzejewski and Olszewski, 1963) showed that yellow-necked mice were commonly aggressive toward the other species, and bank voles withdrew or were chased off by the stronger and heavier species.

Interestingly, bank voles show strategic avoidance of yellow-necked mice through temporal niche switching. Indeed, camera recordings in Carinthia (Austria) showed that, while both species are nocturnal, a high frequency of yellow-necked mice can cause bank voles to invert their activity pattern, becoming diurnal to avoid encounters with yellow-necked mice (Probst and Probst, 2023). A similar temporal partitioning was observed through camera-trapping in a deciduous woodland of southern Tuscany (Italy). Yellow-necked mice were almost exclusively nocturnal, while bank voles showed their phases of activity in the light phase, in particular in the hours after dawn and before dusk (Viviano et al., 2022). However, when this temporal partitioning strategy fails, such as when one species inadvertently invades an active nesting site of the other, aggressive interactions can occur, with potentially dangerous consequences for the voles, which are physically weaker and will likely lose the confrontation. Importantly, notwithstanding the temporal partitioning, it has been observed that there is a small temporal overlap between the activity patterns of bank voles and yellow-necked mice, which comprises a short peri-dawn period and a longer peri-dusk period (Viviano et al., 2022). Interestingly, both our reported cases occurred within this temporal overlap period: the near-lethal case at 7:04 AM (9 minutes after sunrise) and the lethal case at 6:13 PM (78 minutes after sunset).

Furthermore, both bank voles and yellow-necked mice use underground burrows (Ropartz, 1966; Montgomery and Gurnell, 1985; Marchlewska-Koj, 2001; Stradiotto, 2008; Mori and Lazzeri, 2021; Viviano et al., 2022; Górska et al., 2025). It is plausible to imagine that voles occasionally enter the underground nests of yellow-necked mice, and that such invasions, especially in nests with limited escape routes, may result in lethal encounters for the voles. Moreover, it has been reported that, while being capable of burrow-building, yellow-necked mice display a strong tendency to occupy nests that have previously been built by other micromammals, including voles (Grülich, 1980; Amori et al., 1986; Viviano et al., 2022). Rather than digging themselves, they prefer to colonize tunnels and chambers dug by other micromammals. A congener of the yellow-necked mouse, the closely related wood mouse *Apodemus sylvaticus*, shows the same burrow acquisition behavior (Jennings, 1975). For such a behavior, the yellow-necked mouse has been defined “an intruder of existing burrows” (Amori et al., 1986). When a yellow-necked mouse invades a vole nest and finds its occupant(s) inside, it is highly likely that the voles will be either expelled or killed by the mouse. These events may go unnoticed simply due to the cryptic lifestyle of both species, to the location of the killings (intra-burrow), and to the lack of continuous direct observation in natural settings.

Among rodent species, competition-related interspecies killing is rare, as most competition-related interspecies agonistic interactions are just aimed at distancing the competitor. However, it has been previously observed in the herbivorous white-tailed prairie dog (*Cynomys leucurus*), which reduces interspecific competition by killing, without consuming, the herbivorous Wyoming ground squirrels (*Urocitellus elegans*). A long-term study featuring direct observation of a population of free-living white-tailed prairie dogs showed that 47 different individuals killed Wyoming ground squirrels, with 19 individuals being serial killers (Hoogland and Brown, 2016). Interspecies killing was particularly frequent among females, where it was performed by 30% of the individuals. Remarkably, it was seen that, compared to the non-killing females, the killer females had a considerably higher annual and lifetime fitness. Therefore, killing appears to be an extreme way of addressing interspecies competition through permanent removal of the competitor from the game of competition. The lethal and near-lethal attacking that we observed by yellow-necked mice towards bank voles shows the existence of an analogous form of non-predatory, competition-related interspecies killing behavior in the yellow-necked mouse. The reason for which such a behavior could have been selected by natural evolution is that, when the competitor is distanced, it may return. Hence, the advantage for the attacker may be limited in time. On the other hand, if the competitor is killed, this would provide a permanent advantage to the attacker. Moreover, it would exclude the possibility that, after the encounter, the competitor may produce offspring that would become future competitors. Basically, competition-related interspecies killing benefits the attacker by preventing both returns and procreation by the competitor.

Documenting such rare and unexpected behaviors requires targeted, real-time observation, which is still relatively uncommon, especially compared to the entire number of wild animals living in nature. However, such a documentation is fundamental if we wish to describe correctly the general laws that regulate interspecies interactions in nature. Creating theoretical models of interspecies interaction based on extremely partial observations can easily lead to biased interpretations and false deductions. For this reason, it is particularly important to study not only the traces of animal behavior (such as feces, tracks, or nests) but also actual behavioral interactions, using camera traps and other forms of direct monitoring. Our observation chambers are an example of this direct monitoring (Stryjek and Modlinska, 2016; Modlinska and Stryjek, 2016; Stryjek et al., 2018, 2021a, 2021b, 2024a, 2024b, 2026a, 2026b; Parsons et al., 2023a, 2023b; Skorupska et al., 2024). Only through such methods can we uncover behaviors that challenge our expectations, such as territorial killings, infanticide, or unusual forms of social avoidance, offering a fuller picture of behavioral interactions in wild populations.

Detailed behavioral studies of captive rodents have been performed for over 200 years (d’Isa et al., 2024b; d’Isa, 2025b). Studying rodents in a controlled environment such as the laboratory allows a high precision of the observation, removal of any variable which is not of interest and unlimited temporal access to the animals. Nevertheless, often the contexts in which laboratory rodents undergo behavioral tests are quite artificial and, consequently, these tests actually have a scarce ecological and ethological validity. Moreover, some natural behaviors are simply impossible to observe in the laboratory, as the animals do not express them in that environment. On the other hand, studying animal behavior in nature, either by eye or through camera traps, can provide insights of greater ecological and ethological validity. However, both visual observations and camera traps often have the limitation that the target animals are seen from a distance, and fine behavioral interactions not always can be studied with full precision. Instead, our methodological approach, based on the deployment of freely accessible/exitable observation chambers in the natural habitat of the studied animals, allows for a long-term 24/7 monitoring of any behavior that wild-born and free-living animals may display, with a level of visual precision comparable to that of laboratory recordings. Examples of rodent behaviors discovered by our team through this approach are cryoprotective tail-belting (Stryjek et al., 2021a, 2024a, 2026b) and deceptive dodging of a rival (d’Isa et al., 2024a; comment in: Vignieri, 2024). Interestingly, by employing an approach similar to ours, last year the first visually documented case of intraspecies adulticide by the garden dormouse (*Eliomys quercinus*) has been observed (Lang et al., 2025). Basically, the deployment of camera-equipped nestboxes in the natural habitat of the dormice allowed to record an adult garden dormouse that entered a nestbox and killed an adult conspecific that was hibernating inside it. Notably, analogously to our reported case, the corpse of the killed dormouse was not consumed, indicating that this killing was not a form of predation, but rather competition-related. Both our case featuring the yellow-necked mouse and the case featuring the garden mouse, strongly suggest that even in absence of predatory motivations, competition alone can surprisingly lead adult mammals to the killing of other adult mammals. This methodology enabled to highlight in nature the existence of mammalian competition-driven killing of other adult mammals, which is currently a little-known and poorly understood behavior. Indeed, our approach is a particularly promising methodology for the investigation of natural behaviors, especially behavioral interactions that are unexpected, understudied or yet to discover.

## 5. Conclusion

This study presents the first visually documented case of a yellow-necked mouse lethally attacking a bank vole following an intrusion into a nesting site. The event highlights the potential for extreme interspecific aggression in rodents, even among species generally considered non-predatory and primarily omnivorous. While likely such behavior is relatively uncommon, it may currently be underreported due to observational limitations in natural settings. Our findings underscore the value of continuous video monitoring and direct behavioral observation in revealing unexpected or atypical interactions in wild mammal populations. Further research is needed to assess the frequency, triggers, and ecological implications of such lethal encounters among small rodents.

## Ethics Statement

This article is based on the documentation of a fatal attack resulting in evident suffering and visible signs of agony in one of the animals. However, the observations were carried out on free-ranging mice in their natural habitat. The study relied exclusively on video monitoring using food-baited, free-access test chambers, which the animals could enter, exit, or ignore entirely at their own discretion. At no point were the animals captured, restrained, or handled by the researchers. The death occurred as a result of natural interspecific aggression and was not induced by any experimental manipulation or human interference.

According to Polish law (Article 1, Point 2, Act of January 15, 2015, on the Protection of Animals Used for Scientific or Educational Purposes), such non-invasive, observational studies do not require approval from a local ethics committee. All procedures complied with the Polish Animal Protection Act (August 21, 1997), and the study was designed and conducted in accordance with the ARRIVE guidelines (Kilkenny et al., 2010; DOI: 10.1371/journal.pbio.1000412).

## Authors’ contributions

Conceptualization: K.K., R.d.I., P.B., R.S.; Data acquisition: K.K.; Data curation: K.K., R.d.I., M.H.P., P.B., R.S.; Formal analysis: K.K., R.d.I., M.H.P., P.B., R.S.; Methodology: K.K., M.H.P., P.B., R.S.; Visualization: K.K., R.d.I., R.S.; Writing—original draft: K.K., R.d.I., M.H.P., P.B., R.S.; Writing—review and editing: K.K., R.d.I., M.H.P., P.B., R.S. All authors gave final approval for publication and agreed to be held accountable for the work performed therein.

## Supplementary files

Supplementary Video 1: Lethal attack by a yellow-necked mouse on a bank vole.

Supplementary Video 2: Non-agonistic interactions between the killer mouse and a conspecific.

Supplementary Video 3: Yellow-necked mouse burying the body of the dead vole.

Supplementary Video 4: Near-lethal attack by a yellow-necked mouse on a bank vole.

## Footnotes

1 Aggressiveness is the tendency to threaten or actuate an attack (Abramson et al., 2025). On the other hand, dominance is the ability to physically prevail on the opponent. The winner of an agonistic encounter is defined as dominant, while the loser is defined as subordinate. Among free-living rodents, the dominance-subordination relationship is commonly recognized based on which individual makes the other one flee, with the fleer defined as subordinate (Stryjek and Modlinska, 2022; Stryjek et al., 2026a).

2 Also known as striped field mice.

3 Nest-building is an innate behavior that is critical for both self-protection and offspring care (Zichichi Mendis, 2024; Tagawa et al., 2025). Rodent nesting behavior typically follows a specific sequence: a) selection of a location that can offer protection from natural elements and predators; b) searching and collecting nesting material; c) carrying the nesting material to the nesting site; d) sorting: organizing material within the nest, based on the physical characteristics (such as shape and consistency) e) moving and bending the nesting material and positioning pieces of nesting material to form a roundish structure; d) breaking big nesting items into smaller ones; e) using the broken smaller items to form the internal bedding of the structure; f) fluffing: making the central part of the nest more flattened and the border areas of the nest more raised.

4 We identify this yellow-necked mouse as being very likely the same one from the previous day based on morphological similarity and behavioral consistency (persistent nest-building and prolonged time of occupation of the chamber, which was in the order of hours rather than minutes).

5 The phrase “agonistic behavior” was coined and defined by the comparative psychologist John Paul Scott and his collaborators in the early 1950s (Scott and Marston, 1950; Scott and Fredericson, 1951; d’Isa, 2025a). Notably, agonistic behavior does not include just attacks, but also responses such as submission and escape (Scott, 1966). According to the original definition, agonistic behavior is a group of “several patterns of behavior which may provide some measure of adjustment when two organisms come into conflict”, including threat, fighting, freezing, defensive postures and fleeing (Scott and Fredericson, 1951).

